# Molecular regulation of PPARγ/RXRα signaling by the novel cofactor ZFP407

**DOI:** 10.1101/2023.10.25.563939

**Authors:** Alyssa Charrier, Jeremiah Ockunzzi, Siddharth V. Ghanta, David A. Buchner

## Abstract

Cofactors interacting with PPARγ can regulate adipogenesis and adipocyte metabolism by modulating the transcriptional activity and selectivity of PPARγ signaling. ZFP407 was previously demonstrated to regulate PPARγ target genes such as *GLUT4*, and its overexpression improved glucose homeostasis in mice. Here, using a series of molecular assays, including protein-interaction studies, mutagenesis, and ChIP-seq, ZFP407 was found to interact with the PPARγ/RXRα protein complex in the nucleus of adipocytes. Consistent with this observation, ZFP407 DNA binding sites significantly overlapped with PPARγ sites, with more than half of ZFP407 binding sites overlapping with PPARγ DNA binding sites. Transcription factor binding motifs enriched in these overlapping sites included GFY-Staf, ELF1, ETS, ELK1, and ELK4, which regulate key functions within adipocytes. Site-directed mutagenesis of frequent PPARγ phosphorylation or SUMOylation sites did not prevent its regulation by ZFP407, while mutagenesis of ZFP407 regions necessary for RXR and PPARγ binding abrogated any impact of ZFP407 on PPARγ activity. These data suggest that ZFP407 controls the activity of PPARγ, but does so independently of post-translational modifications, likely by direct binding, establishing ZFP407 as a newly identified PPARγ cofactor. In addition, ZFP407 was also found to bind to DNA in regions that did not overlap with PPARγ. These DNA binding sites were more significantly enriched for the transcription factor binding motifs of GFY and ZNF143, which may contribute to the non-PPARγ dependent functions of ZFP407 in adipocytes and other cell types.

## Introduction

More than one-third of adults in the United States are obese [1]. These individuals demonstrate an increased risk for premature mortality given their higher incidence of comorbidities such as type 2 diabetes (T2D) and cardiovascular disease [2]. Although treatments for these conditions exist, safely maintaining a healthy body weight and long-term glucose homeostasis remains a challenge for obese individuals [3]. As key regulatory cells in metabolic signaling and the primary site for lipid storage, adipocytes are central to the pathogenesis of obesity and T2D, displaying promise as targets for therapeutics modulating metabolic processes [4]. Within adipocytes, the peroxisome proliferator-activated receptor (PPAR) family of nuclear receptors, and especially PPARγ, regulate genes involved in glucose homeostasis, adipogenesis, and lipid metabolism [5]. Through an obligate heterodimerization complex with Retinoid X receptors (RXRs), PPARγ recruits additional cofactors necessary to coordinate metabolic processes such as insulin responsive glucose uptake and lipolysis [6].

PPARγ is both necessary and sufficient to regulate adipocyte differentiation and metabolism [7]. PPARγ ligands such as rosiglitazone and pioglitazone, both thiazolidinediones (TZDs), have been utilized as T2D therapeutics on account of their ability to improve insulin responsiveness and lower hyperglycemia, although clinical complications such as increased risk of bone fractures and bladder cancer have limited their use [8]. TZD-induced activation of PPARγ upregulates transcription of genes in key metabolic pathways, such as the GLUT4 glucose transporter, lipoprotein lipase (LPL), and adipocyte fatty acid binding protein (FABP4), among others [9]. In addition to their actions in adipose tissue, TZDs improve insulin resistance in skeletal muscle and liver tissue. However, as PPARγ is more highly expressed in adipocytes, this may be related to endocrine signaling involving free fatty acids and tumor necrosis factor-alpha (TNF-α) originating in adipose tissue [10,11].

Within adipocytes, PPARγ activity displays three primary mechanisms of action. 1) Ligand-independent repression is observed when PPARγ activation inhibits c-Jun N-terminal kinase (JNK) MAPK activity, thereby downregulating gene expression of MAPK targets [12]. 2) PPARγ exhibits both agonist-dependent activation or repression, as in the case of peroxisome proliferator-activated receptor g coactivator 1α (PGC-1α) or nuclear receptor corepressor 1 (NCoR1) biding, respectively [13]. 3) PPARγ activity is also modulated by post-translational modifications such as phosphorylation, acetylation, glycosylation, SUMOylation, and ubiquitination [13]. However, despite decades of study, much of the specific mechanisms underlying PPARγ regulation remains unknown [7,14]. For example, it remains unclear why only a limited number of PPARγ DNA binding sites in adipocytes appear to affect transcription [7,15– 17]. This is likely the result of the combinatorial effects of PPARγ with other uncharacterized transcriptional regulators, which may represent opportunities for novel therapeutics in T2D and obesity treatment by activating subsets of PPARγ targets that improve insulin sensitization without the negative side-effects associated with TZD treatment.

In previous studies, we demonstrated the ability of ZFP407 to positively regulate PPARγ [18]. Utilizing cultured adipocytes, we showed the broad transcriptional effects ZFP407 has on PPARγ signaling, including its regulation of *GLUT4* mRNA, which is directly tied to insulin responsiveness in adipose and muscle tissue [19]. Our *in vivo* studies in mice demonstrated ZFP407 overexpression improves glucose homeostasis [19], while its deficiency causes lipodystrophy and exacerbates insulin resistance [20]. Collectively, our data suggests that ZFP407 is a key regulatory molecule of PPARγ, with unique non-redundant functionality in PPARγ signaling.

Recent studies have demonstrated the impact of context dependence and combinatorial transcription factor action on transcriptional regulation of gene expression [21–26]. With regards to T2D and obesity treatments, this underscores the importance of identifying and characterizing new regulatory cofactors of PPARγ, particularly ones such as ZFP407, which impacts both transcriptional regulatory networks as well as adipocyte development and homeostasis. Utilizing cellular models in this study, we sought to improve our mechanistic understanding of ZFP407’s molecular regulation of PPARγ signaling and better elucidate its role in adipocyte maintenance.

## Materials and Methods

### Materials

Insulin, dexamethasone, and 3-isobutyl-1-methylxanthine, and fetal bovine serum (FBS) were obtained from Sigma Aldrich (USA). Dulbecco’s Modified Eagle’s Medium (DMEM), L-Glutamine/Pen/Strep, 0.05% Trypsin-EDTA, and 0.25% Trypsin-EDTA were obtained from Life Technologies (USA).

### Cell culture

293T cells were cultured in DMEM with 10% FBS and 1x L-Glutamine/Pen/Strep and passaged with 0.05% Trypsin-EDTA. 3T3-L1 cells were cultured, passaged, and differentiated as previously described [27].

### Immunostaining and subcellular fractionization

293T cells were seeded at a density of 5x10^5^ cells onto Poly-D-lysine coated coverslips in 12-well cell culture dishes and incubated overnight at 37°C in 5% CO_2_. Cultured cells were then transfected with either empty vector control, ZFP407 + empty vector control, empty vector control + PPARγ or PPARγ + ZFP407 plasmids using Lipofectamine 3000 (ThermoFisher Scientific, Waltham, MA, USA) according to manufacturer’s protocol. Transfected plasmids encoded PPARγ (Addgene #8862), ZFP407 (cat. #: MR214555, Origene Technologies, USA), or an empty vector control (pRK5-Myc, Clontech, USA). Cells were briefly fixed with 1:1 dilution of methanol and acetone at -20 °C and rinsed with TBS. Double immunofluorescent staining of cell-culture slides for ZFP407 and PPARγ was performed using anti-mouse c-Myc (9E10) (5ug/ml), (Santa Cruz Biotechnology, USA) and anti-rabbit PPAR (D69) (1:100, Cell Signaling Technology, USA) followed by incubation with Alexa-fluor ® 488 goat-anti-mouse (1:1000) and 568 goat-anti rabbit (1:1000) for 1 hour at room temperature. Cell-covered coverslips were mounted onto slides with ProLong™ Diamond Antifade Mountant with DAPI (ThermoFisher Scientific, USA).

Subcellular fractionation was performed using the Nuclear Complex Co-IP kit (Active Motif, USA) according to manufacturer’s protocol until isolation of the nucleus. At this step, the pellet (nuclear fraction) and the supernatant (cytoplasmic fraction) were independently analyzed by western blot.

### Co-IP and western blotting

Co-immunoprecipitation (Co-IP) was performed using the Nuclear Complex Co-IP kit (Active Motif, USA) and Dynabeads™ Protein G for Immunoprecipitation according to manufacturer’s protocol (Invitrogen, USA), and western blotting was performed as previously described [18]. Anti -PPARγ (cat.#: 2430 and 2443), anti-RXRα (cat.#: 3085), and anti-IgG (cat.#: 2729) antibodies were obtained from Cell Signaling (USA). Anti-GAPDH (cat.#: MA5-15738) antibody was obtained from Thermo Fischer Scientific (USA). A custom anti-ZFP407 antibody was generated in rabbits against the C-terminal 149 amino acids of the mouse ZFP407 protein (Proteintech Group, USA), as previously described [19]. Goat anti-rabbit (cat.#: 31460) and goat anti-mouse (cat.#: 31430) secondary antibodies were obtained from Thermo Fisher Scientific (USA). Primary antibodies were diluted 1:1,000, while secondary antibodies were diluted 1:10,000.

### Chromatin Immunoprecipitation (ChIP)-Seq assay and HOMER motif analysis

ChIP was carried out in 3T3-L1 differentiated adipocytes using 30 ug of cell chromatin. ChIP DNA was processed using the Illumina ChIP-Seq kit according to manufacturer’s protocols (Illumina, USA) and sequenced to a depth of 11.5 million reads. The FASTX-Toolkit v0.0.13 was used to quality filter reads using a quality score cutoff of 20. These reads were aligned to the mouse genome (mm9) using Bowtie2 v2.0.6 and any reads with at least one mismatch were discarded. PCR duplicates were also removed using SAMtools v1.3, leaving 9.2 million reads.

The MACS2 algorithm [28] identified 7,313 peaks (q-value < 0.001) using the narrow filter and excluding ENCODE blacklist sites. Control bias-corrected bedGraphs, generated by MACS2 were converted to bigWIGs and used for genome browser visualization. Peaks were segregated based on genomic co-occupancy with either PPARγ or C/EBPα peaks obtained from pervious datasets [29,30] into overlapping and non-overlapping groups. Using these segregated datasets, motif enrichment analysis was performed using Hypergeometric Optimization of Motif EnRichment (HOMER) [31] to determine transcription factor binding motifs enriched in each peak population.

### PPARγ and ZFP407 site-directed mutagenesis

Using the Q5 Site-Directed Mutagenesis Kit according to manufacturer’s protocol (New England Bio Labs, USA), point mutations were created at multiple sites in PPARγ. Lowercase nucleotides indicate mutated residue sites. Serine 112 was mutated to alanine using a forward primer with the sequence 5’- AGAAC CTGCA gctCC ACCTT ATT -3’ and a reverse primer with the sequence 5’- ACTTT GATCG CACTT TGGTA TTC -3’. Serine 112 was mutated to aspartate using a forward primer with the sequence 5’- AGAAC CTGCA gatCC ACCTT ATTAT TC -3’ and a reverse primer with the sequence 5’- ACTTT GATCG CACTT TGG -3’. Lysine 107 was mutated to arginine using a forward primer with the sequence 5’- AAGTG CGATC aaaCG AGTAG AACCT G -3’ and a reverse primer with the sequence 5’- TGGTA TTCTT GGAGC TTC -3’. Serine 273 was mutated to alanine using a forward primer with the sequence 5’- AACGG ACAAA tGCAC CATTT GTC -3’ and a reverse primer with the sequence 5’- GTCTT TCCTG TCAAG ATCG -3’. Additionally, point mutations were induced at multiple sites in ZFP407. The PPARγ binding motif (amino acids 1980-1984) was mutated from LDALL to ALDAL using a forward primer with the sequence 5’-actgg cgTGT GCTGT CACTG AGTTG -3’ and a reverse primer with the sequence 5’- gcatc tgcGG CTGAG GAGTT GTCAG ATG -3’. The coRNR motif (amino acids 2140-2144) was mutated from ISQII to ISQAA using a forward primer with the sequence 5’- GATCT CTCTC Aggcc gctGT AACAG AAGAG CTAGT C -3’ and a reverse primer with the sequence 5’- TCTCC TTCTG ACTCT ACC -3’.

### PPARγ Luciferase Reporter Assay

293T cells were transfected with Lipofectamine 3000 (Life Technologies, USA) according to manufacturer’s protocol (Life Technologies, USA). 3T3-L1 cells were electroporated as previously described [32]. Transfected plasmids encoded PPARγ (Addgene #8862), PPARγ mutants, ZFP407 (cat. #: MR214555, Origene Technologies, USA), or an empty vector control (pRK5-Myc, Clontech, USA), as well as the PPARγ target gene luciferase reporter plasmid (PPRE) (Addgene #1015). The pRL-SV40 plasmid encoding Renilla luciferase was added for normalization. For 293T cells, 490 ng of plasmid DNA, 100 ug of PPRE reporter, and 10 ng of pRL-SV40 were added per reaction. For 3T3-L1 cells, 99μg of plasmid DNA, 10ng PPRE reporter, and 1μg of pRL-SV40 were added per reaction. Relative luciferase activity was measured 24 h post-transfection with the Dual-Glo Luciferase Assay System (Promega, USA).

## Results

### Nuclear co-localization of PPARγ and ZFP407

To determine the subcellular localization of ZFP407, and potential co-localization with PPARγ, 293T cells were transfected with plasmids encoding expression cassettes of either ZFP407, PPARγ, or both together. Immunostaining demonstrated localization of both proteins in the nucleus when either transfected individually or co-transfected, with considerable overlap between the two proteins when transfected together (Fig. 1A). Co-transfection was repeated in differentiated adipocyte 3T3-L1 cells, after which subcellular fractionation was used to separate the nuclear and cytoplasmic components. Western blotting confirmed ZFP407 and PPARγ localization in the nuclear, but not the cytoplasmic, fraction along with RXRα, another protein localized in the nucleus and a known binding partner of PPARγ [2] (Fig. 1B).

**Fig. 1.**
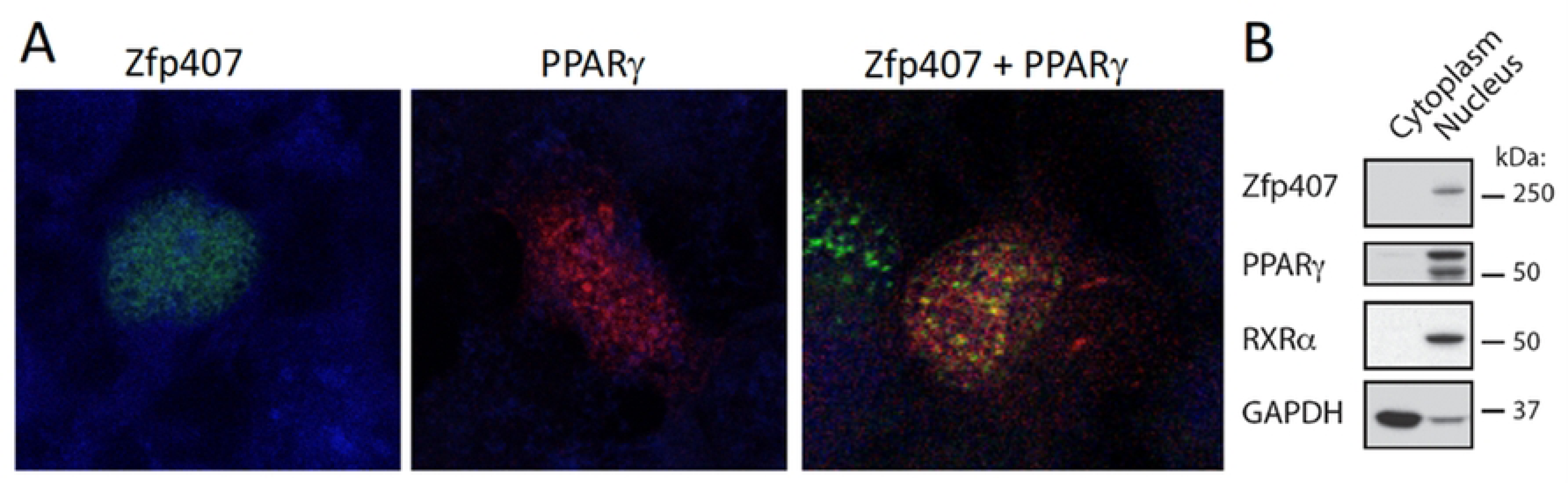
Nuclear localization of ZFP407. (A) Immunostaining of 293T cells co-transfected with plasmids encoding ZFP407 and PPARγ. (B) Subcellular fractionation of 3T3-L1 differentiated adipocytes blotted for ZFP407, PPARγ, and RXRα.

### ZFP407 protein interacts with the PPARγ/RXRα protein complex

Given the overlapping subcellular localizations of PPARγ and ZFP407, we next tested whether these proteins were part of the same complex. ZFP407 and PPARγ were both found to interact with the RXRα protein complex in Co-IP experiments performed on nuclear fractions of 293T cells (Fig. 2A) using an anti-RXRα pulldown of exogenously expressed proteins. Co-IP was similarly performed for the endogenous proteins using the nuclear fractions of 3T3-L1 differentiated adipocytes. ZFP407 was again found to interact with the endogenous PPARγ/RXRα protein complex, showing specific interactions in pulldowns with both an anti-RXRα and an anti-PPARγ antibody (Fig. 2B), indicating ZFP407’s participation in the adipocyte PPARγ/RXRα protein complex.

**Fig. 2.**
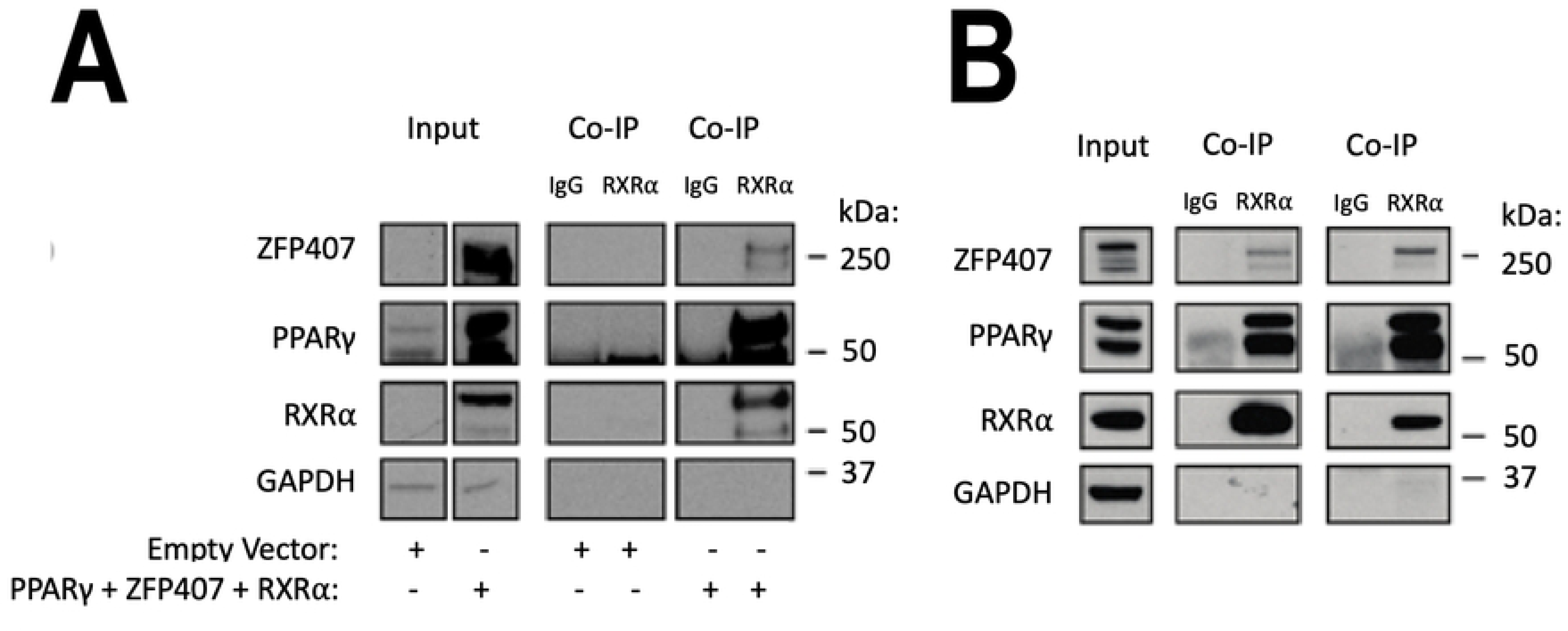
ZFP407 binding in the PPARγ/RXRα protein complex. (A) Co-IP utilizing anti-RXR antibody performed following transfection of either an empty vector or co-transfection of PPARγ, ZFP407, and RXRα plasmids. (B) Co-IP of 3T3-L1 differentiated adipocyte nuclear extracts using matched IgG, anti-RXR, or anti-PPARγ antibodies.

### Overlap between ZFP407 and PPARγ chromatin binding sites

Since ZFP407 is a component of the PPARγ/RXRα protein complex, we hypothesized a significant overlap in chromatin binding sites for these proteins. To test this hypothesis, we performed ChIP-Seq on differentiated 3T3-L1 adipocytes, identifying 7,313 DNA binding sites for ZFP407 throughout the genome. The ZFP407 binding sites identified were aligned with a previous PPARγ ChIP-Seq experiment also performed in 3T3-L1 cells [29]. We discovered that slightly more than half (50.4%) of our identified ZFP407 binding sites overlapped with PPARγ binding sites (Fig. 3A). These included overlapping peaks in the promoter regions of *GLUT4* and *UBQLN1* (Fig. 3B), both genes that have previously been shown to be regulated by PPARγ and ZFP407 individually [18] [33]. When comparing this PPARγ ChIP-Seq dataset [29] with an another published PPARγ ChIP-Seq dataset, also from 3T3-L1 cells [30], we found only 58.7% of sites from the latter ChIP-Seq experiment were identified in both PPARγ ChIP-Seq datasets. This suggests that the 50.4% overlap between ZFP407 and PPARγ in our dataset likely underestimates the degree of overlap when accounting for variance between ChIP-Seq assays examining PPARγ binding in 3T3-L1 cells, as has been noted across multiple studies [30].

**Fig. 3.**
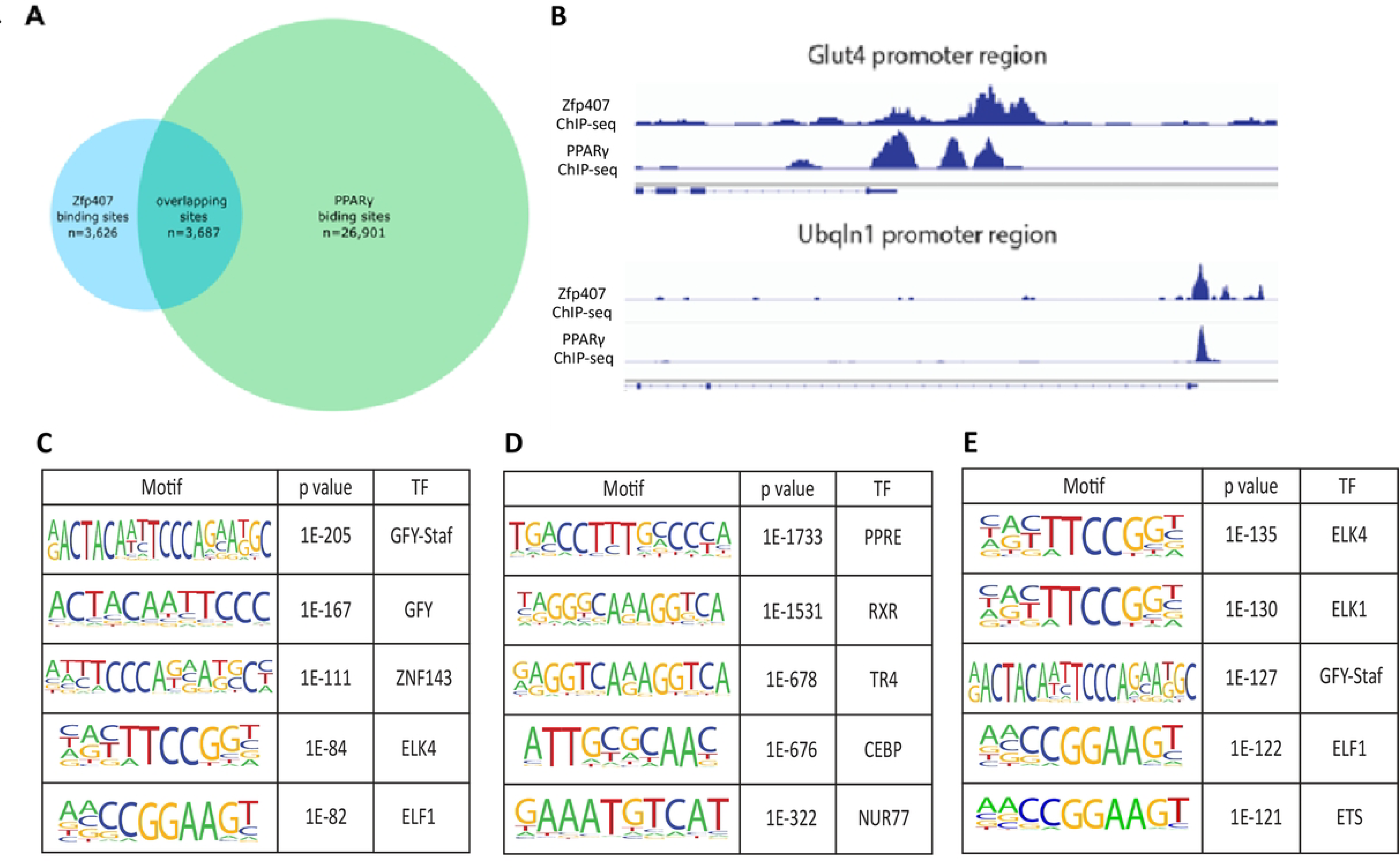
ZFP407 DNA binding sites overlap with PPARγ sites. (A) Overlap of DNA binding sites for ZFP407 and PPARγ determined via ChIP-Seq. (B) Co-occupancy of ZFP407 and PPARγ observed in promoter regions of *GLUT4* and *UBQLN1* genes, PPARγ targets downstream of the Insulin Receptor (INSR). Genomic intron/exon structure for each gene is shown below ChIP data. (C) Top 5 most significant HOMER determined enriched motifs in non-overlapping ZFP407 ChIP-Seq DNA binding sites. (D) Top 5 motifs enriched in non-overlapping PPARγ ChIP-Seq sites. (E) Top 5 motifs enriched in overlapping ChIP-Seq sites. ZFP407 data is from our ChIP-Seq assay, PPARγ data is from Haakonsson et al. (2013).

In addition to PPARγ, we examined the overlap in ZFP407 DNA binding sites with C/EBPα, another key transcriptional regulator in 3T3-L1 cells that is necessary for adipogenesis [34]. In this dataset [29], there was only 36.1% overlap between C/EBPα and PPARγ DNA binding sites and 42.7% overlap between ZFP407 and C/EBPα DNA binding sites. This suggests ZFP407 may more specifically regulate PPARγ signaling than other transcription factors regulating the expression of genes within the canonical adipogenic pathway.

In order to examine any potential non-PPARγ related functions of ZFP407, we segregated the ZFP407 chromatin binding sites into those that overlapped with PPARγ ChIP-Seq peaks and those that did not overlap with PPARγ ChIP-Seq peaks (Supplementary Table 1). The motif analysis program HOMER was used to identify enriched regulatory motifs (Fig. 3C, D, E). Among ZFP407 binding sites that did not overlap with PPARγ, GFY motifs demonstrated the greatest enrichment, with GFY-Staf, ZNF143, ELK4, and ELK1 motifs also demonstrating significant enrichment (Fig. 3C). Among the ZFP407 and PPARγ overlapping sites, ELK4 and ELK1 demonstrated the most significant motif enrichment, with GFY-Staf, ELF1, and ETS also appearing in the 5 most significant motifs (Fig. 3D). The PPAR Response Element (PPRE) was the most enriched motif among PPARγ binding sites that did not overlap with ZFP407, and while for overlapping ZFP407 and PPARγ sites this motif was not among the most significantly enriched motifs, PPRE still demonstrated significant enrichment (p=0.001), consistent with prior results showing ZFP407’s ability to positively regulate PPARγ transcriptional activity towards the PPRE sequence [18].

### PPARγ interaction motifs are required for transcriptional activation by ZFP407

Previous studies have identified consensus protein-protein interaction motifs required for binding to either PPARγ or RXR [35] [36]. The PPARγ binding motif is comprised of an LxxLL amino acid sequence. The RXR binding motif, also referred to as a CoRNR motif, is comprised of an [I/L]xx[I/V]I amino acid sequence. These motifs are both found within the ZFP407 protein, consistent with our co-IP studies showing a direct interaction with the PPARγ/RXR protein complex (Fig. 2A, B). The PPARγ binding motif is located at amino acids 1989-1993 in human ZFP407 and at amino acids 1980-1984 in the mouse ZFP407 (Fig. 4A). The coRNR motif is found at amino acids 2142-2146 in the human ZFP407 and at amino acids 2140-2144 in the mouse ZFP407 (Fig. 4C). Both the PPARγ binding motif and the coRNR motif demonstrate high levels of evolutionary conservation, suggesting an important function for their sequences, and consistent with their function of interacting with the PPARγ/RXR complex. In order to test the function of these two motifs in the transcriptional activity of ZFP407, the PPARγ binding motif was mutated from LDALL to ALDAL, and the coRNR motif was mutated from ISQII to ISQAA, thereby disrupting the binding consensus sites for each. Whereas PPARγ co-transfection with a plasmid encoding wild-type (WT) ZFP407 more than doubled the transcriptional activity of PPARγ (Fig. 4B, D), co-transfection of a plasmid encoding a mutant allele of ZFP407 with either the PPARγ or RXR binding motif completely abolished the transcriptional effect of ZFP407 on PPARγ activity (Fig. 4B, D). This indicates the necessity of both motifs for transcriptional activation of PPARγ by ZFP407.

**Fig. 4.**
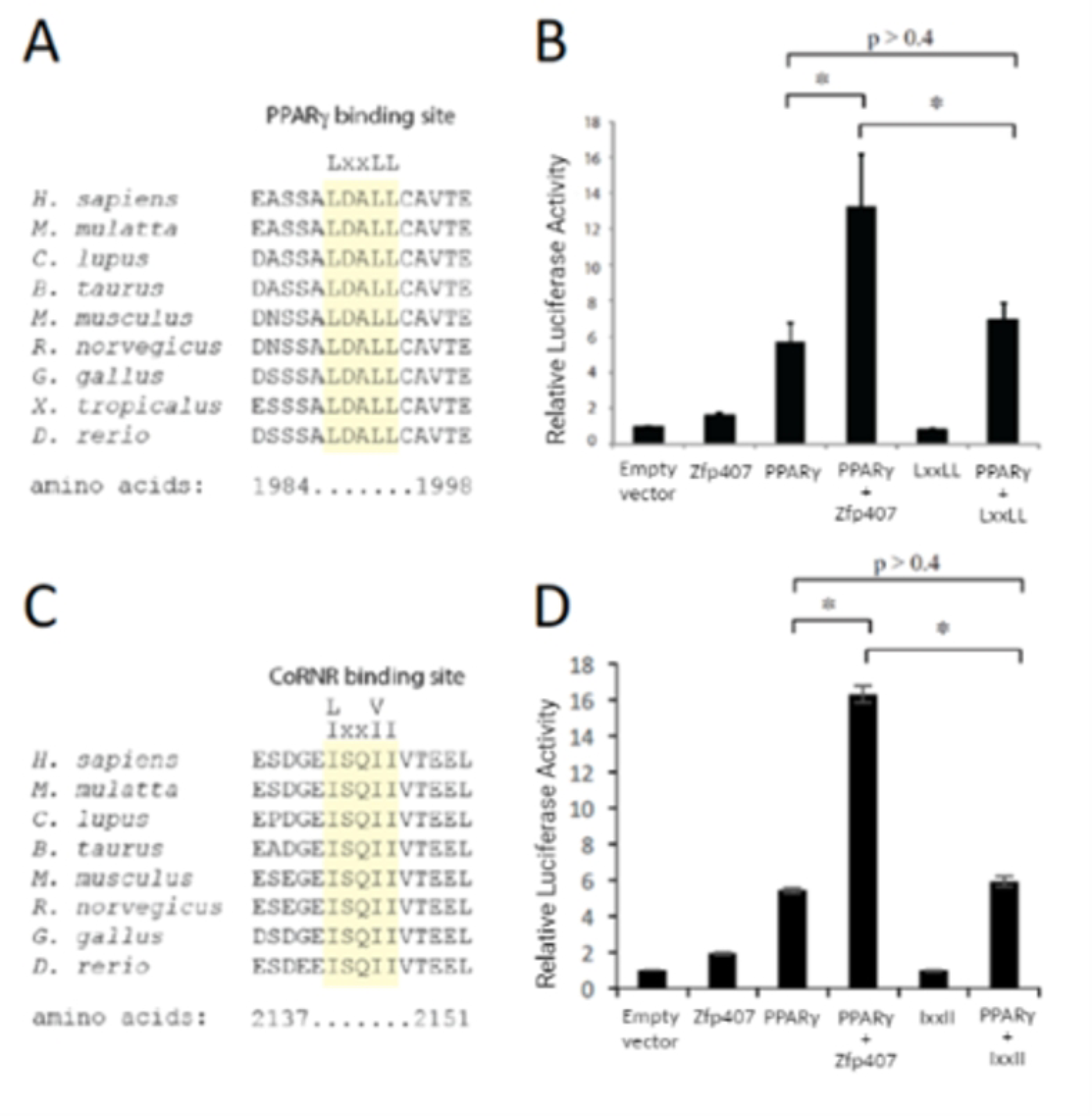
ZFP407 directly binds PPARγ and RXR via LxxLL or [I/L]xx[I/V]I consensus motifs and this interaction is required for transcriptional activation. Putative PPARγ and RXRα interaction motifs in ZFP407. (A) Amino acids 1984-1998 and (C) 2137-2151 of human ZFP407 are shown aligned with other species. Putative binding motifs are highlighted in yellow. (B,D) 293T cells co-transfected with the indicated plasmids and the PPRE PPARγ luciferase reporter. LxxLL and IxxII indicate ZFP407 expression vectors with the indicated motif mutated to contain a binding-disruptive alanine.

### ZFP407 does not modulate PPARγ activity through known PPARγ phosphorylation or SUMOylation sites

Transcriptional activity of PPARγ is moderated by post-translation modifications including both phosphorylation and SUMOylation [37]. Phosphorylation of its serine 112 residue leads to either an increase or decrease of PPARγ activity depending on the context and the protein responsible for phosphorylation [37] [38]. Phosphorylation of PPARγ’s serine 273 reside demonstrated no alteration of PPARγ’s adipogenic induction but altered a subset of PPARγ target genes expression that are commonly dysregulated in obesity including adiponectin and adipsin [39]. Additionally, SUMOylation of PPARγ at lysine 107 is known to decrease its transcriptional activity [40].

To test any potential effect of these modifications on ZFP407’s activation of PPARγ, we created PPARγ mutants with disruptions in each of these post-translational modified sites. The amino acids Serine 112 and Serine 273 were each mutated to alanine to prevent phosphorylation at these sites. Serine 112 was also mutated to aspartate to serve as a phosphomimic. Lysine 107 was mutated to arginine to prevent SUMOylation. When co-transfected with ZFP407, the transcriptional activity of each of these mutant PPARγ alleles increased in a similar manner to that of WT PPARγ, suggesting that the effect of ZFP407 on PPARγ activity is independent of post-translational modifications at these sites (Fig. 5).

**Figure 5.**
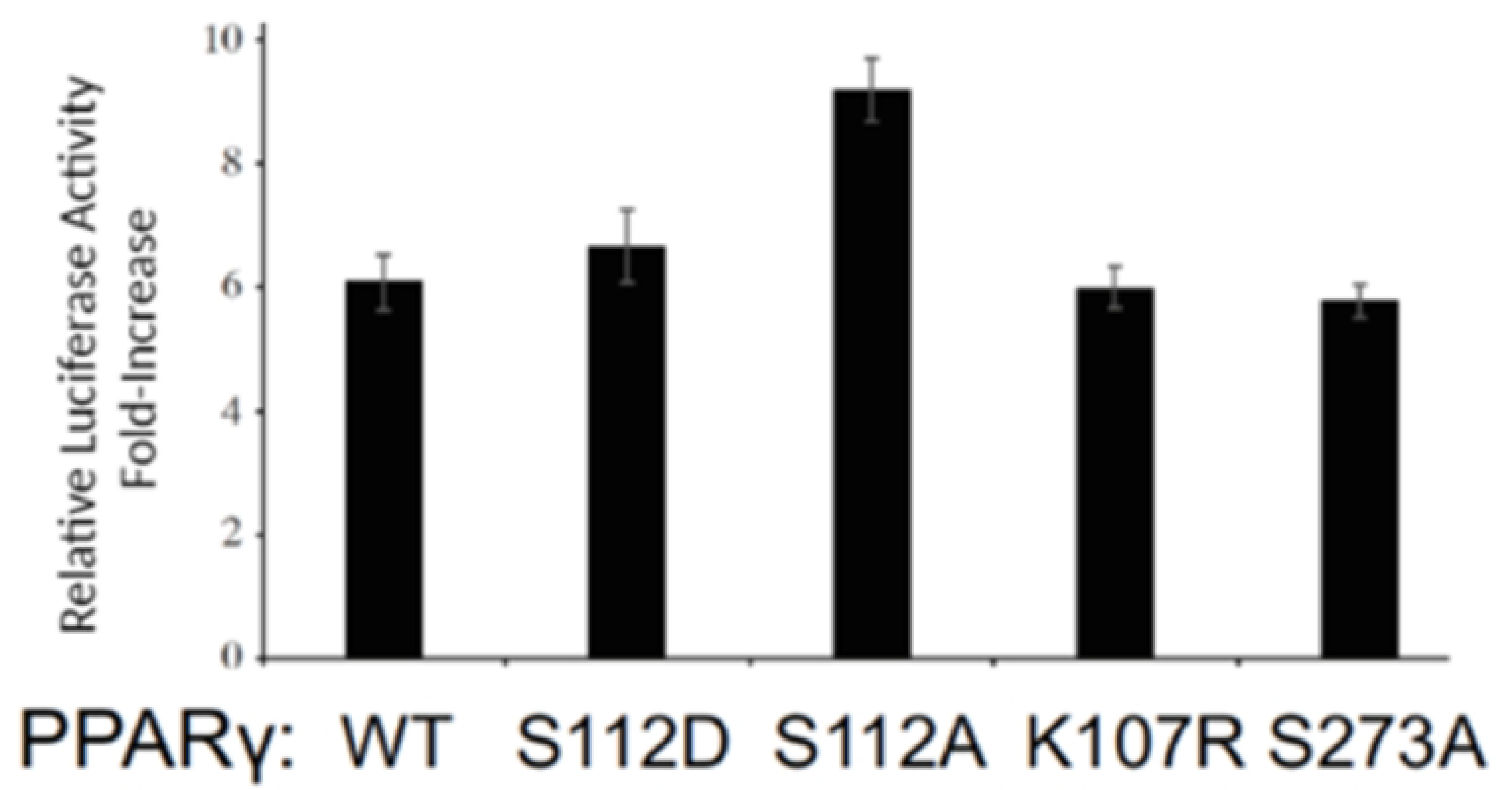
Mutation of PPARg phosphorylation or SUMOylation sites doesn’t prevent its regulation by ZFP407. Co-transfection of differentiated adipocyte 3T3-L1 cells with ZFP407 and wild-type (WT) or indicated PPARγ mutant measured as relative fold-increase of PPRE when compared to an empty vector control.

## Discussion

Immunostaining demonstrated co-localization of PPARγ and ZFP407 proteins in the nucleus (Fig. 1A) when individually or co-transfected. Mutation of common PPARγ phosphorylation or SUMOylation sites did not prevent its regulation by ZFP407 (Fig. 5). In addition, we’ve previously shown that ZFP407 deficiency does not alter PPARγ mRNA or protein levels [18]. These data suggest that ZFP407 controls the activity of PPARγ, but does so independently of post-translational modifications. Consistent with this hypothesis, we demonstrated that ZFP407 is a component of the PPARγ/RXR protein complex, as demonstrated by Co-IP with either an anti-RXR or anti-PPARγ antibody of endogenous proteins from 3T3-L1 adipocytes (Fig. 2B). While it remains unknown if ZFP407 directly binds PPARγ or RXR, or binds indirectly via another cofactor, the presence of ZFP407 in this complex is consistent with a direct effect of ZFP407 on PPARγ signaling.

Cofactors directly interacting with PPARγ often contain at least one LxxLL motif contacting the coactivator binding groove in the ligand-binding domain of PPARγ [41]. ZFP407 contains an evolutionarily conserved LxxLL site (Fig. 4A). When this site was mutated, it negated the effect of ZFP407 on PPARγ transcriptional activity (Fig. 4B). Cofactors that directly interact with RXR often contain a CoRNR motif [I/L]xx[I/V]I [42]. ZFP407 contains an evolutionarily conserved CoRNR site (Fig. 4C). When this site was mutated, it also negated the effect of ZFP407 on PPARγ transcriptional activity (Fig. 4D). Collectively, this data suggests that ZFP407 directly binds PPARγ and RXR via these consensus binding motifs and that this interaction is required for transcriptional activation. The direct binding of ZFP407 could facilitate the interaction between PPARγ and RXR themselves, or facilitate additional cofactor recruitment.

The lone published ChIP-Seq study for ZFP407, a high-throughput analysis of transcription factors in a colon cancer cell line, demonstrated significant DNA-binding overlap between ZFP407 and RXR. The overlap between RXR and ZFP407 ranked in the top 15% of the >7,000 transcription factor pairs analyzed [43]. In addition, we determined that slightly more than half (50.4%) of our ZFP407 ChIP-Seq DNA binding sites overlapped with PPARγ sites from a previously published dataset (Fig. 3A) [29]. These overlapping sites were enriched in GFY-Staf, ELF1, ETS, ELK1, and ELK4 transcription factor (TF)-binding motifs (Fig. 3E). ELK1 has previously been linked to obesity as a target of miRNAs involved in adipogenesis [44], with decreased expression in obese visceral adipose tissue (VAT). Additionally, ELK4 has been linked to adipogenesis, as hyper-methylation of ELK4 binding sites was observed in promoter regions of genes associated with AKT signaling during differentiation of adipose-derived stem cells [45]. *In vitro* studies demonstrated that members of the ETS TF family, and especially ETS2, were necessary for driving early phases of adipogenesis [46]. ETS2 was also found to be enriched in SVF compared to WAT in mice [46]. The enrichment of these motifs suggests ZFP407’s likely role as a cofactor influencing PPARγ’s regulation of the adipogenic program. The remaining top motif hits, ELF1 and GFY-Staf, were enriched in differently methylated regions of mice fed a HFD compared to normal chow [47,48], also suggesting a possible role for ZFP407 in modulating PPARγ function in obese subjects.

ZFP407 binding sites that did not overlap with PPARγ sites were enriched for many similar TF motifs. However, abundance differed most among ZNF143 and GFY motifs, which were more significantly enriched within non-overlapping ZFP407 sites. Within myeloid cells, biding of the ZNF143 TF was shown to be required for C/EBPα transcription [49]. While this finding has not yet been replicated in adipocytes, the abundance of ZNF143 TF biding motifs within our ZFP407 ChIP-seq peaks suggests it may function similarly within adipocytes, where C/EBPα is instrumental during early adipogenic induction [34]. When examining overlap in peaks with another 3T3-L1 ChIP-seq dataset [29], we uncovered 42.7% overlap between ZFP407 and C/EBPα DNA binding sites, further implying a role for ZFP407 in C/EBPα-related regulation of adipocyte function.

Currently, GFY activity within adipocytes has yet to be examined. However, another ChIP-seq performed in 3T3-L1 cells, this time examining TF binding motifs in G protein pathway suppressor 2 (GPS2) promoter regions, observed significant enrichment of GFY sites [50]. This same study demonstrated GPS2 was required for sustaining mitochondrial biogenesis within brown adipose tissue, known for its increased thermogenic energy expenditure [50]. Enrichment of these this motif may indicate that ZFP407, when not cooperating with PPARγ, somehow regulates BAT formation. Previous studies of Zfp407 deficiency in adipocytes, including BAT, demonstrated altered size and morphology of BAT depots [20], morphologically similar to the differences demonstrated by H&E staining of BAT sections in GPS2 adipocyte-KO mice, where increased deposition of lipids was noted [50].

Protein interaction experiments demonstrated binding between ZFP407 and the PPARγ/RXR complex, establishing ZFP407 as a newly identified PPARγ cofactor and suggesting a direct effect on this signaling pathway. With interactions such as direct protein-protein agonism increasingly seen as targets for therapeutic intervention [51], identification of new cofactors, such as ZFP407, in the PPARγ pathway can reveal new drug targets for insulin resistance and T2D. While these studies clearly support the role of ZFP407 as an important cofactor regulating PPARγ/RXRα signaling, they have also provided insight into the other transcriptional pathways potentially regulated by ZFP407. The early embryonic lethality of ZfFP07 knockout mice, which occurs around day e3.5 [20], and is considerably earlier than the mid-embryonic lethality of PPARγ deficient mice by day e10 [52], suggests key functions for ZFP407 beyond the regulation of PPARγ signaling that are required for organismal viability. There remains much to learn regarding the precise mechanism by which ZFP407 controls organismal development and adipocyte function, but these studies have provided new molecular insight into potential pathways and mechanisms by which ZFP407 controls the transcriptional networks that are critical for insulin sensitivity and organismal survival.

---

**Supplementary Table 1. ChIP-seq peaks and motif enrichment hits.** MACS2 identified peaks and motif enrichment as determined by HOMER in ZFP407 DNA binding sites segregated by overlap or non-overlap with PPARγ DNA binding sites.

